# High-dose drug heat map based on organoid array chip for drug selection with high safety and efficacy

**DOI:** 10.1101/2021.05.11.443550

**Authors:** Sang-Yun Lee, Yvonne Teng, Miseol Son, Bosung Ku, Ho Sang Moon, Vinay Tergaonkar, Pierce Kah-Hoe Chow, Dong Woo Lee, Do-Hyun Nam

## Abstract

An organoid array chip was developed by adopting a micropillar and microwell structure to test safety and efficacy of drugs using high dose drug heat map. In the chip, we encapsulated patient-derived cells in alginate and grow them to maturity for more than 7 days to form cancer organoids. When screening drug compounds in a high-density organoid array due to lack of number of patient-derived cells, changing media without damage of organoids is a very tedious and difficult process. Organoids grown in conventional well plates needed too many cells and were also easily damaged due to multiple pipetting during maintenance culture or during experimental procedures. To solve those problem, we applied a micropillar and microwell structure to the organoid array. We used patient-derived cells from patients with Glioblastoma multiforme (GBM), the most common and lethal form of central nervous system cancer, to validate the array chip performance. After forming more than 100µm-diameter organoids in 12 × 36 pillar array chip (25mm × 75mm), we tested 70 drug compounds (6 replicates) with high high-dose to find out high safety and efficacy drug candidates. Comparing the drug response of organoids derived from normal cells and cancer cells, we identified four compounds (Dacomitinib, Cediranib, Ly2835219, BGJ398) as drug candidates without toxicity to GBM cells.

## INTRODUCTION

The 2D monolayer cell culture model has traditionally been used to evaluate the response of cancer cells to different anti-cancer drug compounds. However, when cancer cells are attached to a plastic dish, they grow as a single layer which is morphologically different from the 3D architecture of animal cells grown *in vivo*. The gene expression of the monolayer 2D cells is also different from cells grown in a 3D cell culture model [1]. Additionally, the results of drug screening using a 2D cell culture model are vastly different from those in a 3D culture model [2-6]. As a result, there is great interest and strong motivation by many research groups to improve 3D cell culturing techniques. Generally, 3D cell culture methods can be categorized into scaffold-free methods that allow cells to grow together without an extra-cellular matrix (ECM) and scaffold-dependent methods that cultivate cells with an ECM. Recently, a scaffold-dependent in vitro method was developed to form organoids that recapitulates physiological conditions [7-9]. Those organoids could be used in biomedical research, genomic analysis of various diseases and therapeutic studies [10-13]. In particular, organoids could be powerful tools in drug discovery [14,15] and personalized cancer treatments [9,16].

Growing organoids in a high throughput manner though, is technically difficult due to the potential damage of organoids during pipetting and manipulation. To solve this problem, we applied a micropillar and microwell structure to the organoid array as shown in Fig 1. Previously, we reported the design of a micropillar and microwell chip for culturing 3D cells and using it for drug compound screening tests [17-19]. However, only single cell or small spheroids were exposed to the compounds. In this study, organoid arrays were formed on micropillar chips by growing small spheroids to clumps of cells 100 µm-diameter or larger, as shown in spot images in Fig 1. Culturing organoids this way prevent damage to the organoids from direct pipetting during cell maintenance as the micropillar chip can be directly transferred to a new microwell filled with fresh media. We used the organoid array to evaluate the efficacy and toxicity of different compounds on brain cells derived from patients with Glioblastoma multiforme (GBM), the most common and lethal form of central nervous system cancer. GBM patient-derived cells were supplied from Samsung Medical Center Biobank and their genetic profiles well matched with original tissue in the previous our research [20,21]. After forming more than 100 µm-diameter organoids in 12 × 36 pillar array chip (25mm × 75mm), we tested 70 drug compounds (6 replicates each) to evaluate individual drug efficacy and toxicity in high dose. With limited number of patient derived cells (PDC), the purpose of this paper is the primary screening to select efficacy drugs among many drugs without risk of brain cell cytotoxicity. Therefore, we choose a high-dose of 20 µM to exclude cytotoxicity compound in high-dose. Under this condition, we selected high efficacy drug candidates without cytotoxicity risk by evaluating drug response of organoids derived from normal cells (astrocyte) and cancer cells (GBM PDC).

**Figure 1.**
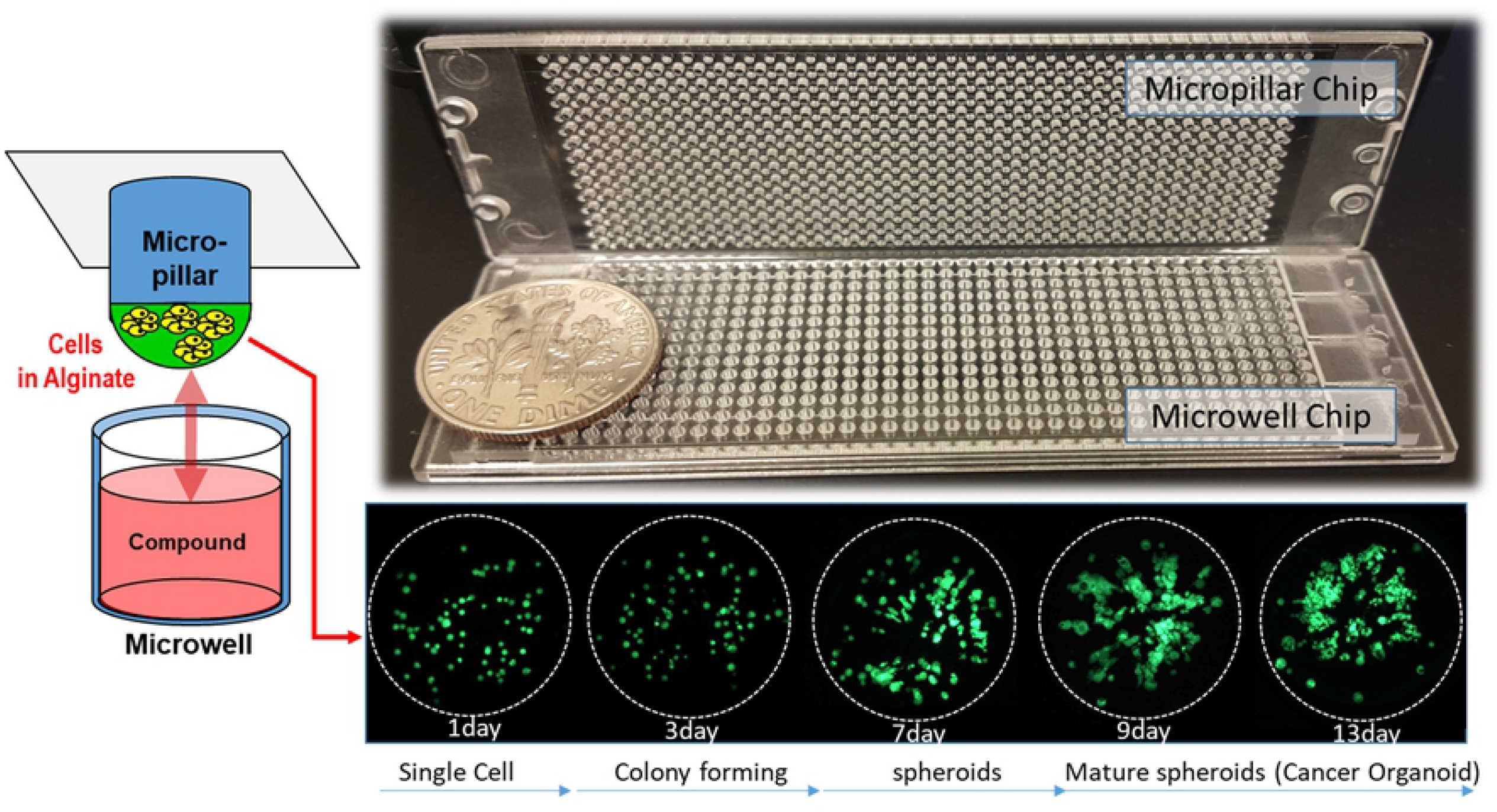
Cancer organoid array chip designed based on micropillar and microwell chips. Green represents living cells and organoids more than100 µm in diameter, which were formed after 7 days.

## MATERIALS AND METHODS

### Experimental procedures

Approximately 100 cells (patient-derived GBM cells and astrocytes) in 50 nL with 0.5 % (w/w) alginate were automatically dispensed onto a micropillar chip by using ASFA™ Spotter ST (Medical & Bio Decision, South Korea). The ASFA™ Spotter ST uses a solenoid valve (The Lee Company, USA) for dispensing 50 nL droplets of the cell–alginate mixture and 1 µL of media or compounds. After dispensing the cells and media in micropillar and microwell respectively (Fig 2A), the micropillar chip containing human cells in alginate was combined (or “stamped”) with the microwell chip filled with 1 µL the fresh media (Fig 2B). The micropillar and microwell chip in the combined form are shown in (Fig 2b). After incubation for 3 days at 37°C, cells started to form spheres. We replaced the media with fresh media every 2-3 days and allowed the cells to continue growing for up to 7 days or when the size of spheres become larger than 100 µm. Micropillar chips were spaced 200 µm from the microwell chip to enable uniform CO_2_ penetration into the organoids (Fig 2C). An incubation chamber was used while incubating the cells on the chip to prevent media evaporation (Fig 2D). At day 7, drug compounds were added to cancer organoids (Fig 2E). In Fig 2F, cell viability was measured with Calcein AM live cell staining dye (4 mM stock from Invitrogen), which stains viable cells with green fluorescence. The fabricated micropillar/microwell chips were compared with five 96-well plates in (Fig 2G). A fully grown cancer organoid with and without addition of drug compounds was shown in (Fig 2H).

**Figure 2.**
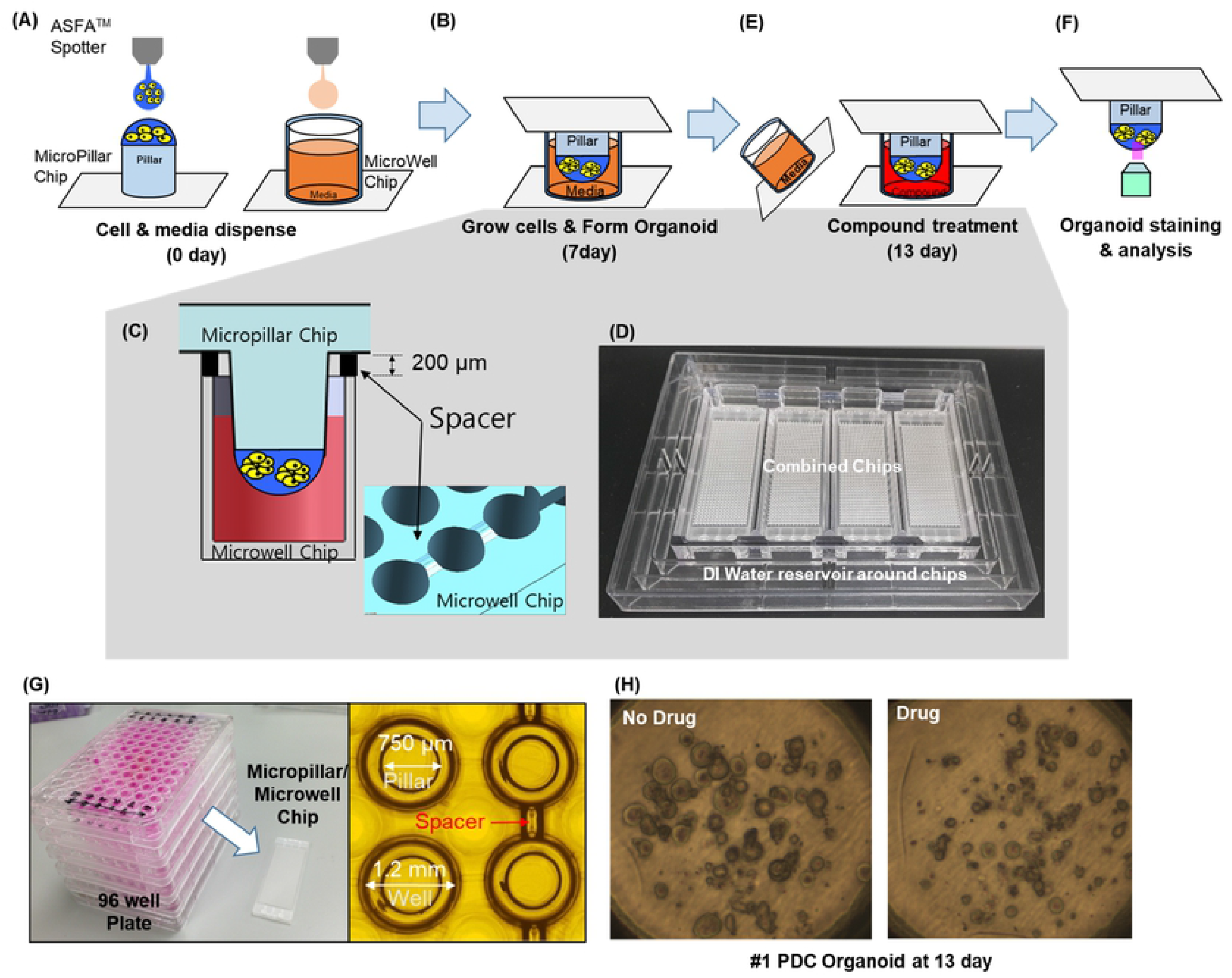
Experimental procedure and fabricated chips. (A) Cells and media are dispensed on micropillar and in microwell, respectively. (B) Cells were immobilized in alginate on the top of the micropillars and dipped in the microwells containing growth media for 7 days to form mature spheroids (organoids). (C) Spacer on the microwell chip created gaps between pillar and well chips for CO_2_ gas exchange during long term cell culture. (D) Chip incubation chambers designed to prevent media evaporation during long term cell culture. (E) Compounds are dispensed into the microwells and Organoids are exposed to the compounds by moving the micropillar chip to a new microwell chip. (F) Organoids are stained with Calcein AM, and the dried alginate spot on the micropillar chip is scanned for data analysis. (G) photo of the combined micropillar and microwell chip compared with conventional 96 well plate. (H) Cancer organoids images with and without compound.

### Fabrication of micropillar/microwell chips and incubation chamber for organoid array

Previously, the micropillar/microwell chip platform was used for short term cell culture to form spheroids. When we applied this to long term organoid cell culture, we found that the low volume of media (1 µL) in each microwell evaporated quickly and there was also a lack of CO_2_ supply in the tightly packed chips. To solve these technical issues, the micropillar/microwell chips were modified. When we tightly sealed the micropillar and the microwell chip to prevent evaporation, there was insufficient CO_2_ in the microwell. We then created a gap between the micropillar and the microwell chips to allow CO_2_ to easily flow into the wells. The modified micropillar and microwell chips were manufactured by plastic injection molding. The micropillar chip is made of poly (styrene-*co*-maleic anhydride) (PS-MA) and contains 532 micropillars (0.75 mm pillar diameter and 1.5 mm pillar-to-pillar distance). PS-MA provides a reactive functionality to covalently attach poly-L-lysine (PLL), ultimately attaching alginate spots by ionic interactions. Plastic molding was performed with an injection molder (Sodic Plustech Inc., USA).

The incubation chamber for the micropillar/microwell chips (Fig 2D) was fabricated by cutting cyclic olefin copolymer (COC) with a computer numerical control (CNC) machine. COC was selected because of its high transparency, excellent biocompatibility, and adequate stiffness for physical machining. As shown in (Fig 2D), reservoir around 4 combined chips was filled with deionized (DI) water to prevent evaporation. We checked that 5.3% of media evaporated in the incubation chamber during a period of 13 days.

### Astrocyte and Patient-derived cell culture

We purchased NHA-astrocyte AGM (LONZA, Cat. No:cc-2565) and cultured it with ABM Basal media (LONZA, Cat. No:cc-3187) that was added with AGM SingleQuot Kit Suppl.&Growth Factors (LONZA, Cat. No:cc-4123). Patient-derived GBM cells were obtained from GBM patients who underwent brain tumor removal surgery at the Samsung Medical Center (Seoul, Korea). Informed consent was obtained from all patients. Following a previously reported procedure [13], surgical samples were enzymatically dissociated into single cells. Four patient-derived cells were obtained from four GBM patients. Dissociated GBM cells were cultured in cell-culture flasks (from Eppendorf, T-75) filled with Neurobasal A (NBA) conditioned media. The NBA conditioned media comprised N2 and B27 supplements (0.53 each; Invitrogen) and human recombinant bFGF and EGF (25 ng/ml each; R&D Systems), hereafter, referred as NBE condition media. Cell flasks were placed in a humidified 5% CO_2_ incubator (Sheldon Mfg., Inc.) at 37°C. The cells were routinely passed every 4 days at 70% confluence. For the experiment, cell suspensions were collected in a 50-ml falcon tube from the culture flask. GBM cells were then suspended in 5 mL of NBE condition media. After centrifugation at 2000 rpm for 3 min, the supernatant was removed, and the cells were re-suspended with NBA conditioned media to a final concentration of 10 × 10^6^ cells/mL. The number of cells in the NBA conditioned media was calculated with the AccuChip automatic cell counting kit (Digital Bio, Inc). The rest of the cells were seeded at a concentration of 2 × 10^6^ cells in a T-75 flask containing 15 mL of NBA conditioned media.

### Efficacy/toxicity test

To validate organoid array chip, seventy compounds were evaluated using four PDCs and normal astrocytes (Normal brain cell) for drug efficacy and toxicity test, respectively. The compounds, which showed low cytotoxicity against astrocytes and high efficacy against four PDCs, were selected as the best candidates for GBM cancer. For experimental testing, we selected seventy compounds whose targets are well-known including: epidermal growth factor receptor (EGFR), phosphoinositide 3-kinase (PI3k), mechanistic target of rapamycin (mTOR), vascular endothelial growth factor (VEGFR), c-mesenchymal-epithelial transition factor (c-Met), and fibroblast growth factor receptors (FGFR). These compounds are in phase III or phase IV trials or are approved oncology drugs by the US Food and Drug Administration (FDA). With limited number of Patient-derived cells, the purpose of this paper is the primary screening to select efficacy drugs among many drugs without risk of brain cell cytotoxicity. Therefore, we choose a high dose of 20 µM based on TMZ, one of the most used drugs for GBM [22].

Figure 3 shows the layout of 72 different compounds in the scanning image of the micropillar chip. Each compound has 6 replicates. The top row (compound 1, 37) is the DMSO control without compound treatment. The enlarged scanning images show live organoids (green lumps) treated with different compounds. Empty (block) circle means organoids were not viable due to effects of the compounds.

**Figure 3.**
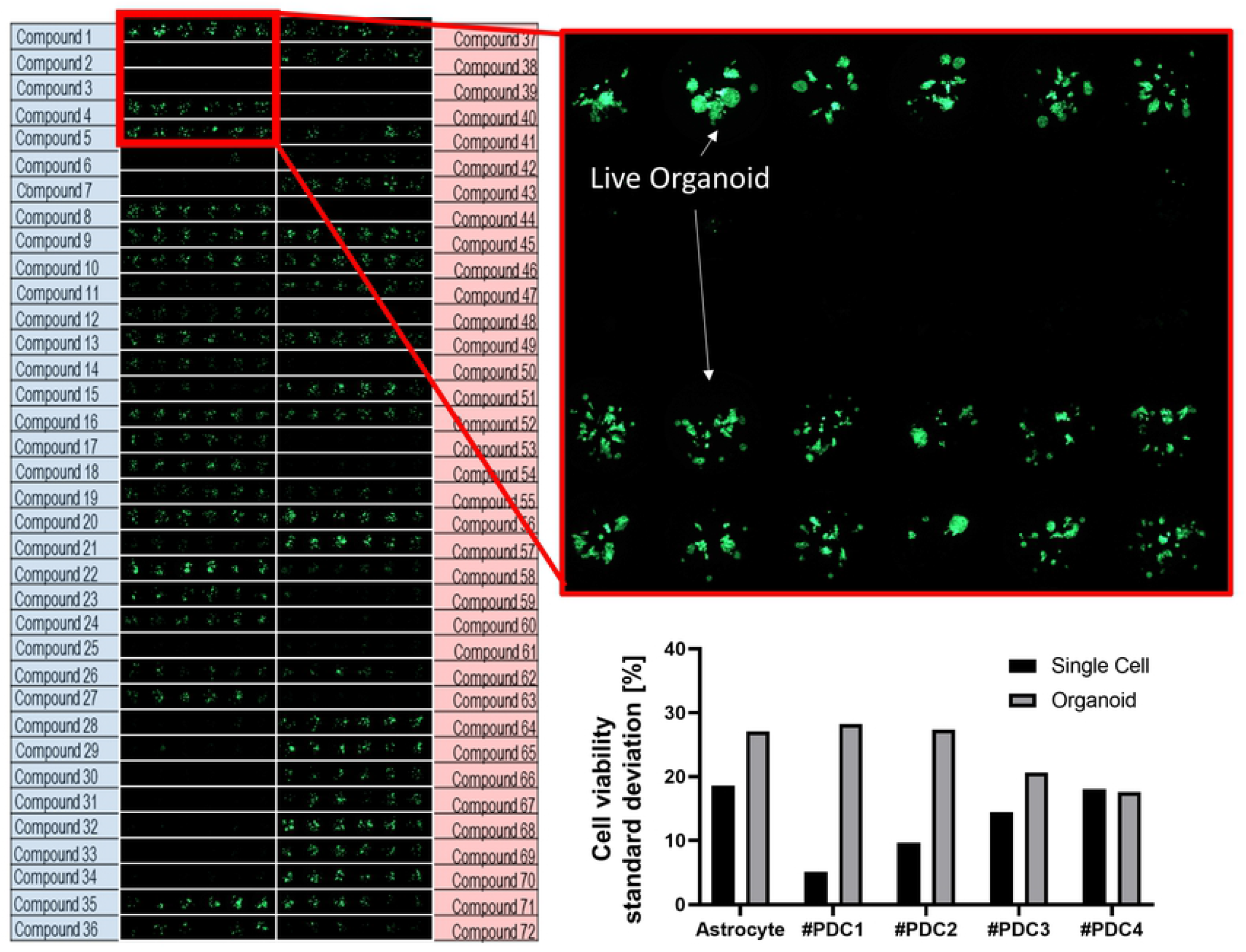
Scanning image of micropillar chip exposed with 72 (including 2 DMSO) compounds. The enlarged image shows staining of viable cell organoids (6 replicates). The graph shows the standard deviation of cell viability in condition of no compound (control) at single cell model (7 day culture) and organoid model (13 day culture)

We compared the response of compounds to single cells and organoids separately. Single cells were treated at Day 1 to the different compounds and then stained at Day 7. For organoids, compounds were added at Day 7 (after cells form organoids) and then stained at Day 13, as shown in (Fig 2). In typical drug screening, drugs were exposed for twice the cell doubling time. In case of GBM patient derived cells, the doubling time is an approximately 3 to 4 day. Thus, we expose the compound for 7 days.

### Viability measurement of organoid

Live organoids were stained using Calcein AM (4 mM stock from Invitrogen). The staining dye solution was prepared by adding 1.0 µL of Calcein AM in 8 mL staining buffer (MBD-STA50, Medical & Bio Device, South Korea). To measure organoid viability quantitatively after staining the alginate spots, organoids on the micropillar chip were scanned. As shown in (Fig 3), scanned images were obtained with an automatic optical fluorescence scanner (ASFA™ Scanner ST, Medical & Bio Device, South Korea). From the image of the organoid spots, the total green-fluorescent intensity from live organoids in each spot was extracted using Chip Analyzer (Medical & Bio Device, South Korea). The software extracted total green-fluorescent intensities (8 bit Green code among RGB code: 0∼255) from living organoids in each cell spot on the scanned chip by setting up an analysis boundary (the circles in Fig 4). The relative organoid viabilities were calculated by dividing green intensities of compounds with one of the controls (no compound treatment).

**Figure 4.**
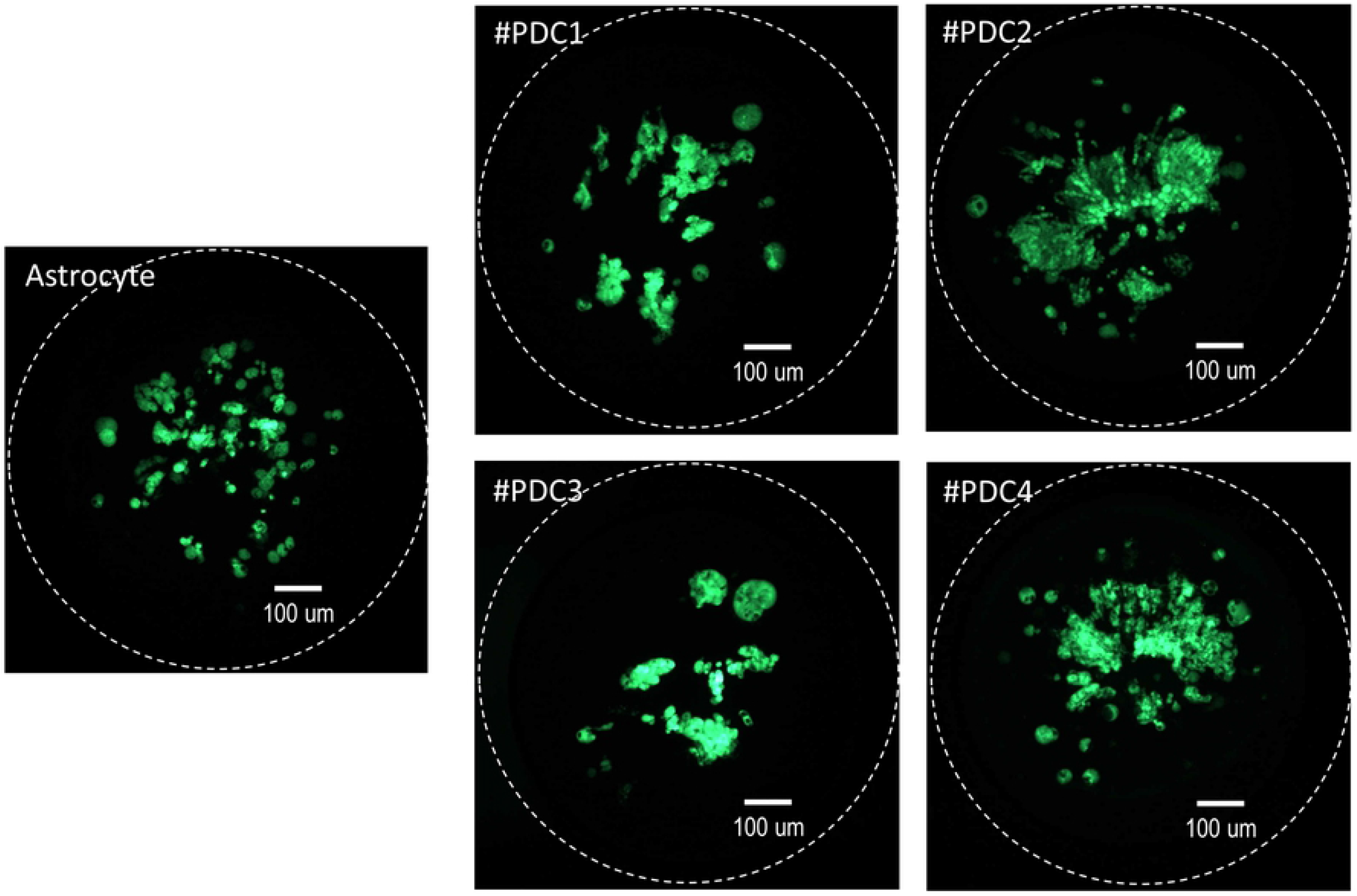
Organoid image of astrocyte (brain normal cell) and four GBM patient derived cells after 13 days of culturing

### Statistical analysis

All data were expressed as the mean±standard deviation (SD) of three or more independent measurements. Student t-test as well as χ 2 test were used to test the statistical significance using Microsoft Excel 2010. P-values < 0.05 were considered statistically significant.

### Ethical statement

This investigation was conducted in accordance with the ethical standards of the Declaration of Helsinki and national and international guidelines and was approved by the Institutional Review Board at Samsung Medical Center in Seoul, Korea (IRB No. 201512092)

## RESULTS and DISCUSSION

Previously, the efficacy and cytotoxicity of 70 compounds were screened in micropillar/microwell chip platform [18]. Compared to Temozolomide (TMZ), one of the most popular drugs for GBM, we found that cediranib showed high drug efficacy to GBM patient derived cells without toxicity. However, many compounds including TMZ showed cytotoxicity against normal glial cell (astrocytes) because of strong drug toxicity towards single cells when the drug was added before the cells formed colonies. In this paper, we modified micropillar/microwell chip platform for long-term cell culture to form cancer organoids. Spacer gaps were introduced to allow CO_2_ exchange and incubation chambers were designed to prevent evaporation of media. We used the array to grow cancer organoids for 7 days, after which the organoids were exposed to 70 compounds for another 7 days. Drug efficacy and cytotoxicity were measured. As shown in (Fig 4), normal astrocytes and four GBP patient-derived cells formed mature spheroids (or organoids) whose size are more than 100 µm-diameter.

### Toxicity test (astrocytes) with Organoids

As shown in (Fig 5), most compounds show high cytotoxicity when the compounds were exposed to single cells for 7 days. Most compounds had similar cytotoxicity when used on single cells or organoid astrocytes. However, fifteen drug compounds demonstrated a low cytotoxicity to organoids (mature spheroid) or astrocytes (viability is higher than 50%). Of these fifteen drug compounds, 6 compounds showed low cytotoxicity on single cells while 9 compounds showed low cytotoxicity in both single cells and organoids, as shown in (Fig 6). In previous research, TMZ which is widely known for its low cytotoxicity, exhibited high toxicity (with astrocyte viability < 16%) after treating the cells for 7 days but showed low toxicity (with astrocyte viability > 90%) after a 3-day treatment period. Previous studies assessed the cytotoxicity of TMZ using a 3-day treatment period only [23]. Using astrocyte organoids, TMZ showed no cytotoxicity even though it was treated for 7-days. TMZ is commonly used to treat GBM cancer patients. Using TMZ drug testing results on organoids of astrocyte as a benchmark, more low cytotoxicity compounds were found such as Dacomitinib, Axitinib, LEE011, LY2835219, BGJ398, LGK-974, PHA-665752, Amoral and Raloxifene.

**Figure 5.**
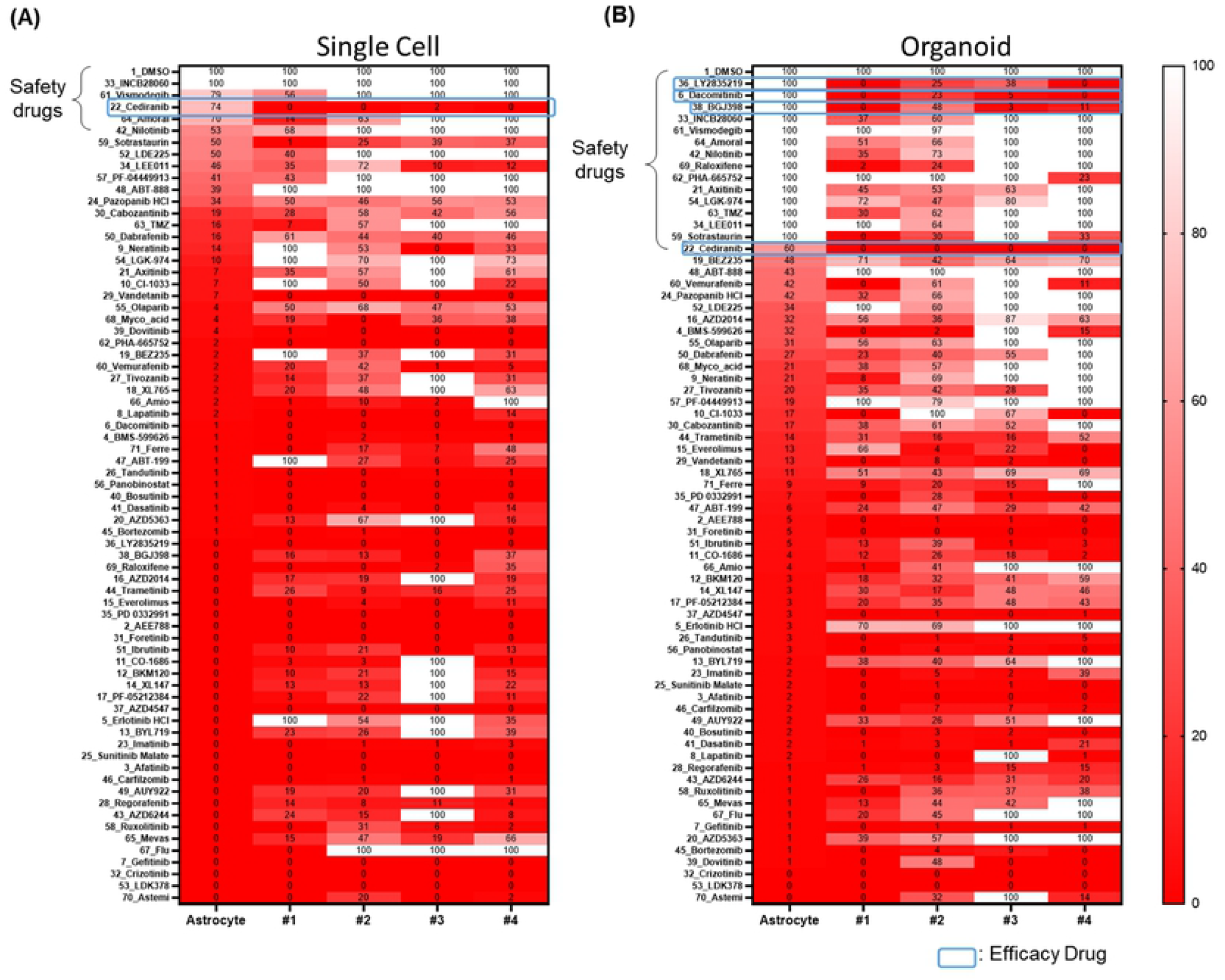
Relative cell viabilities when single cells or organoids were exposed to drugs for 7 days. Cell viability was measured using the green fluorescence intensity of living cells. The relative cell viabilities were normalized based on the cell viabilities of DMSO only condition. Green color shading means no cytotoxicity due to more than 50% astrocyte viability. (A) Single cell condition. (B) organoid condition

**Figure 6.**
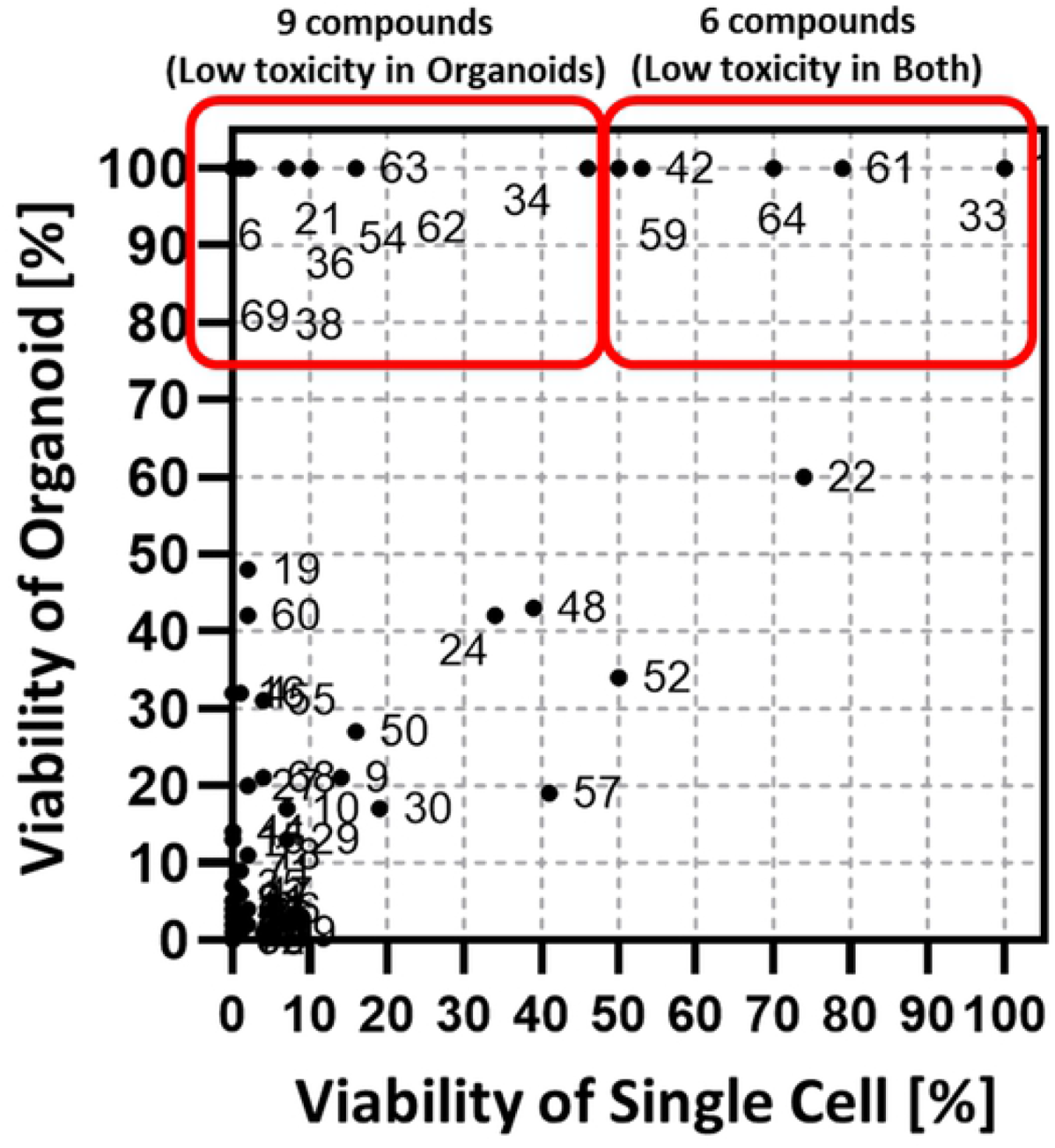
The cell viabilities of single cell and organoid of astrocyte exposed with 70 compounds for 7 days.

### Efficacy test of 70 compounds (four patient derived GBM cells) with organoids

The cell viabilities of normal astrocyte and four patient derived GBM cells treated with different drug compounds, either in single cell or organoid conditions were shown in (Fig 5). Most of the targeted compounds exhibited high efficacy with 20 µM dosages in both single cell and organoid conditions. In previous work [18] where single cells were exposed to compounds for three days, cediranib exhibited high efficacy in all four patient-derived GBM cells and exhibited no cytotoxicity towards astrocytes. In seven-day compound treatment of single cells, most compounds showed high cytotoxicity in astrocytes as well as high efficacy in four patient-derived GBM cells. Six compounds showed low cytotoxicity (viability>50%) but only cediranib and Sotrastaurin demonstrated high efficacy to four-patient-derived GBM cells as shown in (Fig 5). However, in organoid culture, fifteen compounds showed low cytotoxicity in astrocytes and four compounds (Dacomitinib, Cediranib, Ly2835219, BGJ398) showed high efficacy in four patient-derived cells (Fig 7). As shown in (Fig 3), replication is six for each compound. The graph in (Fig 3) shows standard deviation of cell viabilities in single cell and organoid in no compound condition (control). Single cell models showed under 20% standard deviation and organoid model under 30% standard deviation. Big organoids showed higher variation. However, the efficacy drugs (Dacomitinib, Cediranib, Ly2835219, BGJ398 selected in (Fig 7) have P-value of less than 0.05. Compared to viability of no drug (control), organoid viabilities of four compounds were found to be statistically significant low and four compounds highly efficient in GBM cells. Dacomitinib, Ly2835219, and BGJ398 showed high cytotoxicity when compounds were treated early in single cell culture before forming spheroids of astrocytes. However, mature spheroids (organoids) of astrocytes displayed high resistance to those three compounds while four patient-derived GBM cells had markedly reduced viability to those three compounds. In other research papers, those three compounds showed the high efficacy to GMM cells or patients. Abemaciclib (LY2835219), a drug that inhibits Cyclin-Dependent Kinases 4/6 and crosses the Blood-Brain Barrier, demonstrates *in vivo* activity against intracranial human brain tumor xenografts [24]. In preclinical test, dacomitinib showed its effectiveness against glioblastoma [25]. BGJ398 is currently in Phase II clinical trials to assess anti-tumor efficacy in recurrent GBM patients (*NCT number:* NCT01975701). Our study showed that by employing our high-throughput organoid array model, we identified new drug candidates such as Dacomitinib, Ly2835219, BGJ398, and cediranib for GBM. This serves as a proof-of-concept study to identify new drug candidates for other cancers using our high-throughput organoid array chip.

**Figure 7.**
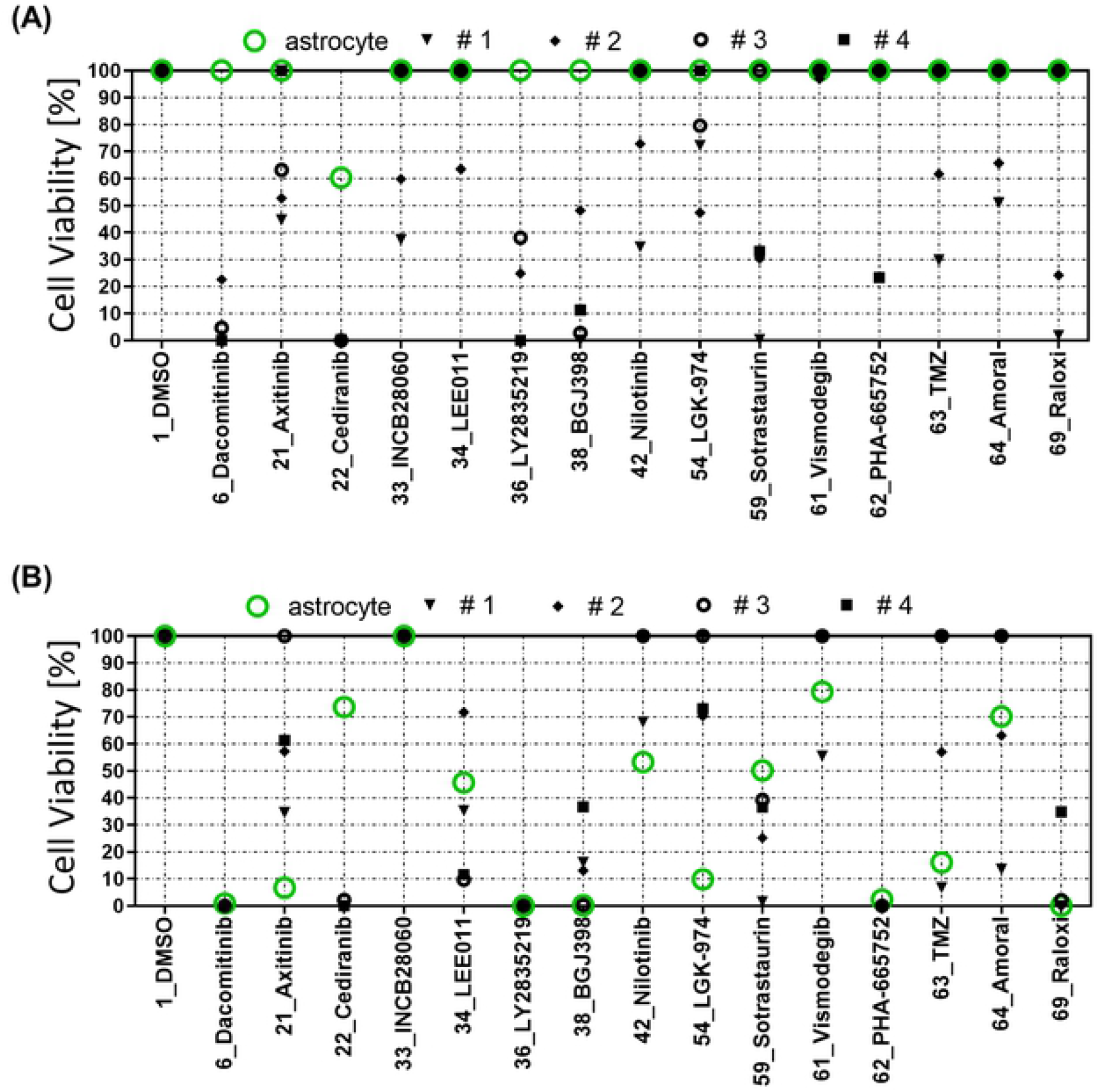
Compound heat map of organoids and single cells. (A) In organoid, cells were cultured for 7 days to form organoids (spheroid dimeter is over 100 µm) and organoids were treated with compounds for a further 7 days. (B) In single cell culture, compounds were exposed to single cell 1 day after seeding cell and single cells were treated with compounds for 7 more days.

## CONCLUSION

Cancer organoid array chip was developed using the micropillar and microwell structure to be used for drug screening. In the chip, we encapsulated cells in alginate and grow patient derived cells for more than 7 days to form cancer organoids. Importantly, this method also prevents accidental damage to organoids during cell maintenance and culturing. We used cells derived from patients with GBM, the most common and lethal form of central nervous system cancer, to validate the organoid array chip performance. After forming more than 100 µm-diameter organoids in 12 × 36 pillar array chip (25 mm × 75 mm), we treated the organoids with 70 different drug compounds (6 replicates each) to evaluate drug compound efficacy. We identified new drug candidates with high safety and efficacy for GBM using this high-dose drug heat map array.

## ACKNOWLEDGMENTS

Declaration of Conflicting Interests The authors declare no potential conflict of interests with respect to the research, authorship, and/or publication of this article. This research was supported by the National Research Foundation of Korea(NRF) grant funded by the Korea govern-ment(MSIT) (No. 2020R1I1A3066550), the Korea Medical Device Development Fund grant funded by the Korea government (the Ministry of Science and ICT, the Ministry of Trade, Industry and Energy, the Ministry of Health & Welfare, the Ministry of Food and Drug Safety) (NTIS Number: 202012E15-04).

We thank PuRPOSE Programme Study Team for their helpful discussions and technical support on this project.

